# A spectral analysis approach to detect actively translated open reading frames in high-resolution ribosome profiling data

**DOI:** 10.1101/031625

**Authors:** Lorenzo Calviello, Neelanjan Mukherjee, Emanuel Wyler, Henrik Zauber, Antje Hirsekorn, Matthias Selbach, Markus Landthaler, Benedikt Obermayer, Uwe Ohler

## Abstract

RNA sequencing protocols allow for quantifying gene expression regulation at each individual step, from transcription to protein synthesis. Ribosome Profiling (Ribo-seq) maps the positions of translating ribosomes over the entire transcriptome. Despite its great potential, a rigorous statistical approach to identify translated regions by means of the characteristic three-nucleotide periodicity of Ribo-seq data is not yet available. To fill this gap, we developed RiboTaper, which quantifies the significance of periodic Ribo-seq reads via spectral analysis methods.

We applied RiboTaper on newly generated, deep Ribo-seq data in HEK293 cells, to derive an extensive map of translation that covers Open Reading Frame (ORF) annotations for more than 11,000 protein-coding genes. We also find distinct ribosomal signatures for several hundred detected upstream ORFs and ORFs in annotated non-coding genes (ncORFs). Mass spectrometry data confirms that RiboTaper achieves excellent coverage of the cellular proteome and validates dozens of novel peptide products. Collectively, RiboTaper (available at https://ohlerlab.mdc-berlin.de/software/) is a powerful method for comprehensive *de novo* identification of actively used ORFs in the human genome.

## INTRODUCTION

Ribosome Profiling (Ribo-seq) is a method to globally investigate protein synthesis^1^ by sequencing RNA fragments protected by engaged ribosomes. Mapping these ribosomal footprints to the transcriptome produces a detailed quantitative picture of translation across thousands of genes^2^. In the few years since its first description, Ribo-seq has been applied to monitor translation in different organisms^3^ and biological conditions^4^. Applications cover a wide range of subjects, including the identification of alternative start codons^5^ and the definition of short peptides^6,7^ as well as quantitative aspects such as translation rates or codon efficiency^8^.

With an increasing number of annotated non-coding RNAs, a central question is whether some of these transcripts contain short translated ORFs missed by annotation efforts. To this end, a number of studies proposed *ad-hoc* Ribo-seq based global metrics that aimed at defining the translational status of such non-coding transcripts^9–11^. The mere presence of Ribo-seq reads in regions of the transcriptome does however not imply the presence of actively elongating ribosomes. The mode of Ribo-seq read length distributions typically falls at *^~^*29–30nt, which is the known fragment size protected by 80S ribosomes^12^. Consequently, this subset of ribosomal footprints displays a striking bias towards the translated frame^1,6,9^, which can be used to infer, for each read, the position of the peptidyl-site (P-site) compartment of the translating ribosome^9^. This sub-codon resolution holds great promise to discriminate between ribosomal coverage and a periodic footprint profile (PFP), i.e., a consistent, 3-periodic codon-by-codon alignment pattern across a transcript. Yet, this property has only recently been exploited within summary statistics to identify small translated ORFs^6,13^ or to explore translation on multiple frames^14^.

Despite the recent development Ribo-seq analysis tools^15–17^, a statistically rigorous method using this property to identify translated regions has not been proposed. Here we present a computational approach, based on spectral analysis, to comprehensively identify the set of PFPs in a given Ribo-seq sample. Our method, called RiboTaper, exploits the sub-codon resolution of Ribo-seq reads to call high-confidence translated loci, and reconstructs the full set of ORFs in annotated coding and non-coding transcripts. We quantify how RiboTaper outperforms alternative approaches, show that a limited number of non-coding RNAs contain actively translated ORFs, analyze evolutionary signatures in different ORF classes, and cross-reference the identified ORFs with proteome-wide experimental evidence.

## RESULTS

### The RiboTaper strategy for testing Ribo-seq sub-codon profiles

The main idea of our approach is to provide a statistical test to identify PFPs, indicative of consistent codon-by-codon ribosomal movement across a putative translated region. This test is the central part of the RiboTaper method (Fig. 1a): We first define P-site positions for the majority of mapped reads, according to their aggregate profiles over annotated start codons^6,9^ (Fig. 1a, Supplementary Fig. 1). Next, we create data tracks for every annotated exonic region, and use the multitaper approach^18^ to test for significant PFPs in exonic P-site profiles (Fig. 1a). Finally, exonic profiles are merged according to the annotated transcript structures to detect translation on *de novo* identified ORFs (cf. Methods, Software availability, Supplementary Software).

**Fig. 1:**
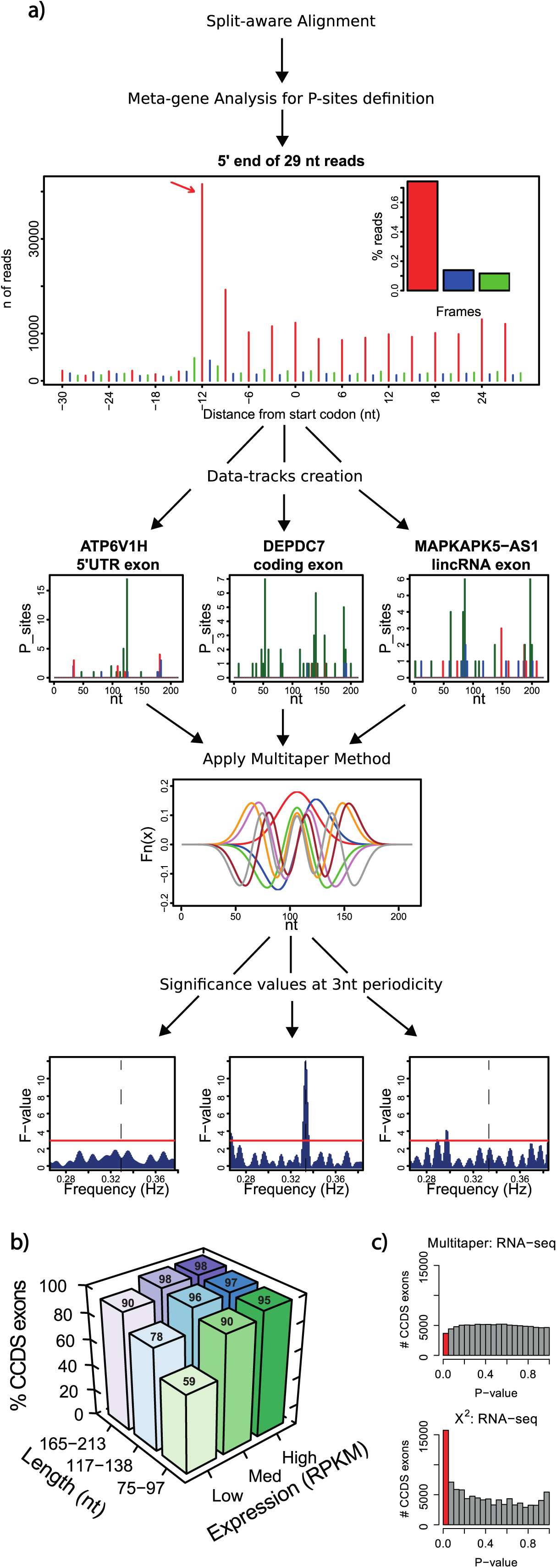
The RiboTaper workflow. Ribo-seq reads are first mapped to the genome using a split-aware aligner (e.g. STAR). a) To infer P-site positions, aggregate profiles are created over annotated start and stop codons, for different read lengths. Data tracks (P-sites tracks shown) are created for all annotated exons; three examples for a coding exon, UTR exon and exon in a non-coding gene are shown. The tracks are analyzed with the multitaper method, which applies a set of orthonormal functions (known as Slepian sequences, shown for 7 tapers) to allow for robust estimates of every frequency component. Finally a statistical significance test is performed for each exon at a frequency of 3nt. Three examples of length- and coverage-matched exons are derived from different transcript classes and exhibit differences in significant 3 nt periodicity. b) Impact of ORF length and expression on the percent of expressed CCDS exons detected to exhibit significant 3nt periodicity (exact value on each bar). Low, Med, and High bins correspond to 2.34–3.84, 5.91–9.14, and 15.08–29.77 RPKMs respectively. c) Histogram of p-values for all CCDS exons tested applying MultiTaper (top) and Chi-squared (bottom) tests to RNA-seq data as a control that should not exhibit extensive periodicity. P-values < 0.05 are shown in red.

The multitaper approach applies a set of orthogonal window functions (tapers) to a discrete signal before computing its Fourier transform. The transformed windowed outputs are averaged to obtain a smoothed spectrum amenable to a non-parametric test for detecting significant frequencies^19,20^ (cf. Methods). In this way, the multitaper method tests directly for the presence of PFPs, in contrast to current approaches that look at enrichment of Ribo-seq reads over RNA-seq (Translation Efficiency^1^) or a preference for reads to align to one frame.

### RiboTaper defines active translation with high sensitivity and specificity

To assess the performance and utility of RiboTaper, we generated a deep ribosome profiling dataset in HEK293 cells following established protocols (cf. Methods). We obtained >29 Mio reads, out of which >25 Mio aligned to the genome (Supplementary Table 1). Profiles at consensus coding exons (CCDS annotation, see Methods) displayed a striking frame preference, in excellent agreement with annotated transcript types and regions (Supplementary Fig. 2).

To evaluate the sensitivity of the multitaper, we applied it on profiles from CCDS exons of different length and Ribo-seq coverage (Supplementary Fig. 2–4). As expected, the test achieved higher sensitivity on longer exons and benefitted from higher coverage (Fig 1b). Twenty-four tapers exhibited the best combination of sensitivity and specificity (Supplementary Fig. 5).

We then benchmarked the multitaper against a Chi-squared significance test as baseline. This test uses a null hypotheses of a uniform frame P-site distribution and corresponds to the basic assumption behind the ORFscore^6^. However, sequencing protocols are affected by different sources of variability that cause non-uniform distribution of reads^21^. Further, insufficient depth may also lead to sampling biases and spurious enrichments. To evaluate specificity on an appropriate negative sample, i.e. regions without 3nt-periodic reads, we applied the tests on RNA-seq data of annotated CCDS exons.

At a significance level of 0.05, the multitaper displayed slightly lower sensitivity when compared to the Chi-squared test (87% vs. 94% of positive exons, Supplementary Fig. 2–4). However, when applied to RNA-seq profiles, the Chi-squared test showed a worrisome number of positive calls compared to the multitaper test (16% vs. 3.5%, Fig. 1c, Supplementary Fig 2–4). We observed strongly skewed p-values for the Chi-squared test but a desirable near-uniform distribution of the multitaper p-values on the RNA-seq (Fig 1c). This high specificity is critical when exploring translation outside of protein-coding regions and directly pertains to Ribo-seq data: Integrated in our analysis pipeline, the Chi-square detected 67 translated ORFs in snoRNAs and snRNAs - transcripts unlikely to generate peptides -- while the multitaper detected only 3. A principled statistical framework eliminates the need for heuristic parameters or ad-hoc cutoffs, a crucial step when applying score-based metrics on different datasets (Supplementary Fig. 6).

### Full ORF reconstruction captures known and novel translated ORFs

We next created transcript tracks for *de novo* translated ORF identification, using 3nt periodicity and frame definition by the P-site positions (Fig 2a, Methods). We classified ORFs based on annotation categories and genomic position relative to known coding regions (Fig. 2b-c, Supplementary Fig. 7, Methods). This led to ^~^21,000 translated ORFs in ^~^14,000 expressed genes, in coding and non-coding transcripts, across a wide range of expression values (Fig. 3a).

**Fig. 2:**
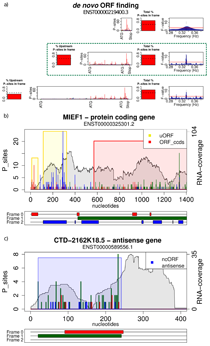
*De novo* ORF reconstruction and examples of significant ORFs. In case of multiple possible start codons it is necessary to discriminate between internal methionines and initiation codons. a) For a given transcript we anchor the analysis on the stop codon and utilize the most upstream in-frame ATG with more than 5 P-sites positions (>50% in-frame) between it and its closest downstream ATG (green dashed box). Examples of translated ORFs are shown in b) for two uORFs and one CCDS ORF for MIEF1 and in c) for ncORF in CTD-2162K18.5. For a given transcript each plot depicts the RNA-seq coverage (grey), the potential ORFs of the three possible frames (red, green, and blue), and the ORFS with significant PFP colored by the annotation category. All of the PFP ORFs detected are supported by peptides from mass-spec data except for the most 5’ uORF in MIEF1.

The vast majority of ORFs in protein-coding genes overlapped known CCDS coding regions; 369 non-CCDS protein-coding genes were identified as harboring translated ORFs ("nonccds_coding_ORFs"). We detected >600 genes with translated upstream ORFs (uORFs) and 54 genes with downstream ORFs (dORFs; cf. Methods). We additionally identified ORFs in 504 non-coding genes (ncORFs). These ORFs belong mainly to pseudogene, antisense, and lincRNA biotypes (Fig. 3b).

**Fig. 3:**
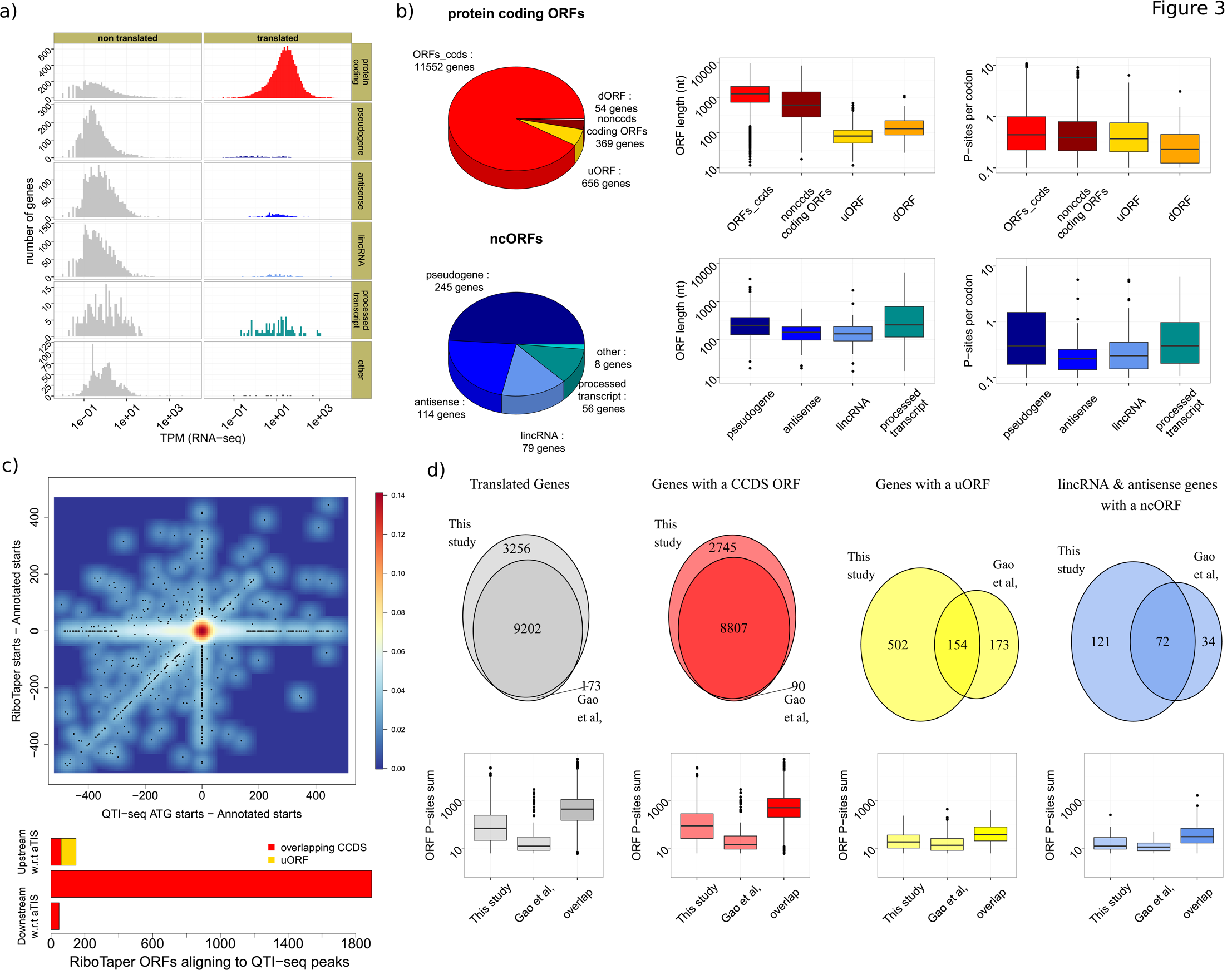
Comparative analysis of translated ORFs from coding and non-coding regions. a) Histogram of expression levels for genes with (right) and without (left) translated ORFs by annotation category. b) Number, length and coverage of protein coding ORFs and ncORFs, along the different sub-categories. c) Scatterplot (top) of the distance between reported QTI-seq ATG peaks and annotated start codons (x-axis) vs distance between RiboTaper ORFs starts and the annotation (y-axis). Barplot (bottom) of the number of start positions identified by both QTI-seq and RiboTaper with respect to the annotated translation initiation site (aTIS). d) Overlap (top) and coverage (bottom) of ORFs identified in the Gao *et al* data set compared to our data set, split by ORF category.

RiboTaper identified ORFs exhibited a protein-coding like distribution of Ribo-seq read lengths as quantified by the FLOSS score^11^, across all ORF categories (Supplementary Fig. 8). The reconstructed ORF coordinates agreed with translation initiation sites defined by QTI-seq^22^ or the reference annotation (Fig. 3c). Compared with the reference, 149 upstream initiation sites were detected by both QTI-seq and RiboTaper, mostly corresponding to uORF start codons (Fig. 3c). 52 internal starts were identified by both QTI-seq and our method. Approximately 1000 QTI-seq ATG start codon candidates did not overlap with either annotated or RiboTaper-defined start codons. Using Ribo-seq data from the same study, more than 99% of Ribotaper-identified CCDS ORFs were also found in our data (Fig. 3d). Agreement dropped to 68% for lincRNAs/antisense ORFs and 47% for uORF-containing genes, possibly due to their relatively short length and low expression levels (Fig. 3d).

To demonstrate the general applicability of RiboTaper, we identified thousands of coding ORFs and ncORFs in Ribo-seq data of the zebrafish embryo^6^ (Supplementary Fig. 4, Supplementary Table 2–3, Supplementary File 1). Among the identified ncORFs was the recently discovered ORF in the lincRNA *toddler*^23^, which encodes a small polypeptide morphogen essential for zebrafish embryonic development (Supplementary Fig. 9).

### Conservation patterns underline diverse roles of open reading frames in different RNA classes

To gain insights into the identified ORFs in categories outside of annotated coding regions, we analyzed evolutionary conservation patterns of ORFs in different categories. CCDS ORFs and non-CCDS coding ORFs exhibited high nucleotide conservation^24^, followed by pseudogenes and processed transcripts (Supplementary Fig. 10). Different from *bona fide* coding regions, we observed elevated nucleotide conservation only around start and stop codons for uORFs and processed transcripts (Fig. 4a).

**Fig. 4:**
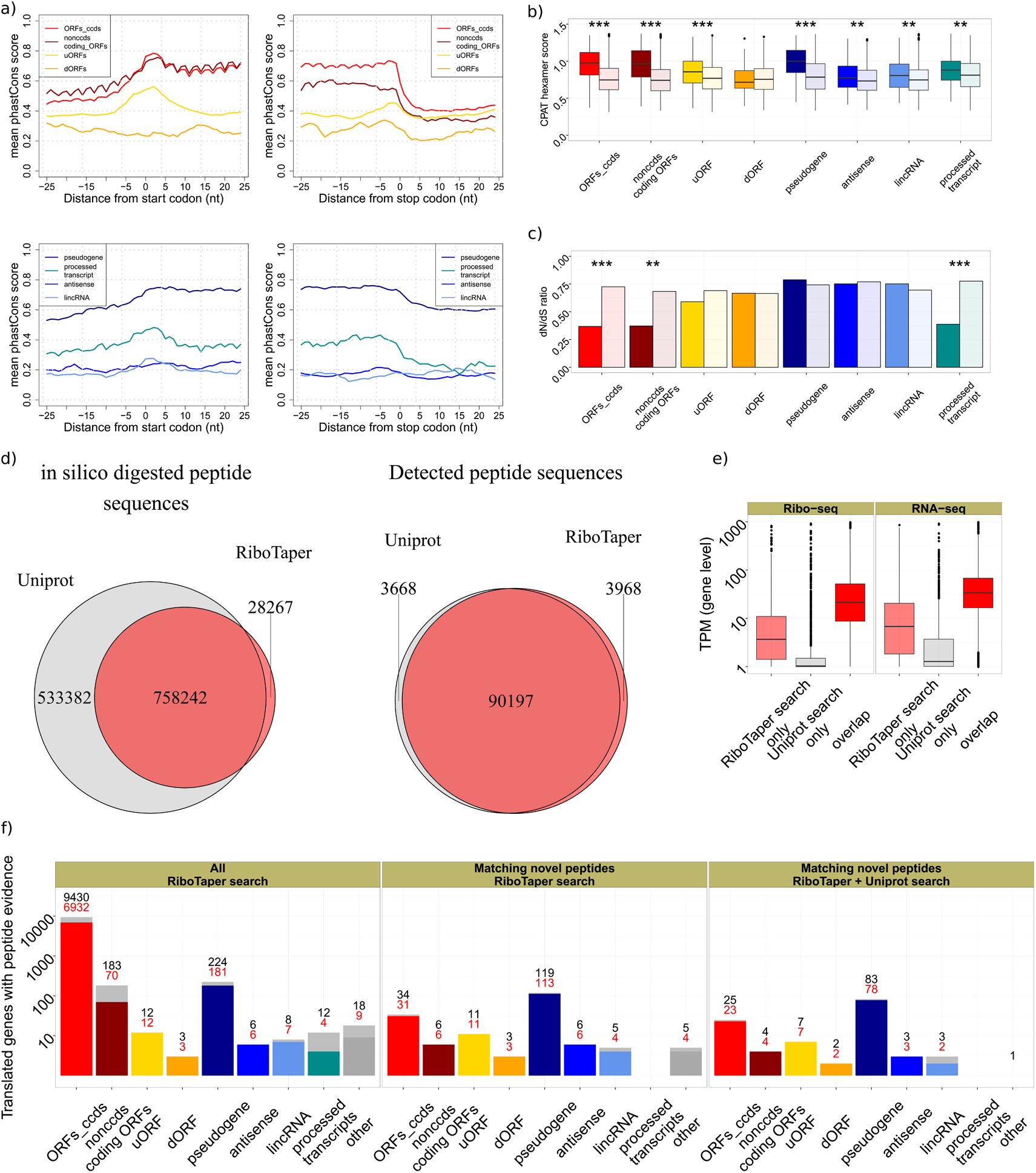
Conserved and non-conserved RiboTaper-identified ORFs define the cellular proteome. a) Local nucleotide conservation (computed by PhastCons) in a +/- 25 nt window around start and stop codons positions. b) Comparison of length- and conservation-matched control ORFs to the different ORF categories, by CPAT hexamer score and c) dN/dS ratio. ORFs_ccds and nonccds coding ORFs <300 nucleotides long were used in this analysis, to match negative control ORFs (Supplementary Fig. 10). d) Overlap between the protein databases derived from Uniprot or RiboTaper ORFs for all possible peptide sequences (left) and for detected peptide sequences (right). e) Gene expression levels (Ribo-seq and RNA-seq) for genes showing peptide support in the two search strategies (RiboTaper vs Uniprot). f) The number of genes containing at least one RiboTaper identified ORF with peptide evidence by ORF category (left). Genes containing at least one RiboTaper identified ORF with novel peptide evidence (not found in Uniprot, human entries, rel. October 2014) using the RiboTaper database (middle; 191 ORFs in 189 genes) or a database of the union of RiboTaper and Uniprot entries to exclude potential cross-matches (right; 157 novel peptides mapping to 129 ORFs in 127 genes). Red numbers indicate evidence from uniquely assigned peptides.

Next, we assessed whether sequence conservation reflected evolutionary selection on the encoded protein sequence, and whether this selection persists in the human population. We quantified coding potential by means of hexamer sequence statistics^25^ and tested the preference for synonymous vs. non-synonymous SNPs in the human population using appropriate length- and conservation-matched controls(Fig. 4c, Supplementary Fig. 10, Methods). For CCDS and non-CCDS coding, ORFs, as well as processed_transcript ncORFs, nucleotide conservation was accompanied by good hexamer scores and a depletion of nonsynonymous SNPs (dN/dS). uORFs also showed conservation on the nucleotide level but no significant enrichment of synonymous substitutions (dN/dS), suggesting potential regulatory rather than protein-coding roles, since their position but not sequence is conserved.

Additional ncORFs categories showed low nucleotide conservation (except for pseudogenes), a positive trend for hexamer scores, but no depletion for nonsynonymous SNPs. Codon substitution patterns across vertebrate species^26^ led to a similar outcome (Supplementary Fig. 10). Taken together, ncORFs detected by RiboTaper do not necessarily entail conservation and selection patterns similar to protein-coding regions.

### Ribosome profiling serves as an effective proxy to define the cellular proteome

The ORFs identified by RiboTaper covered a wide range of expression values (cf Fig. 3a), to an extent that its coverage might exceed deep mass spectrometry datasets in defining the cellular proteome. To evaluate this, we created a custom database from the set of identified ORFs to match the spectra of a recent HEK293 tandem mass spectrometry dataset^27^. The RiboTaper peptide set corresponded to ~59% of the peptides in Uniprot (human entries, rel. October 2014), and an additional 2% of non-Uniprot candidates. The RiboTaper set matched >90.000 peptide sequences, belonging to >8.000 genes (Fig. 4d, Supplementary Fig. 11), similar to the results of the full Uniprot search.

Over 3900 peptide sequences were found only by our custom search but not using the Uniprot database (1% FDR, Supplementary Table 4). In turn, our search missed a similar number of peptides. RiboTaper-only peptides matched more spectra than UniProt-only peptides, despite being shorter and with lower matching scores (Supplementary Fig. 11). We found little evidence for expression or translation for most of the Uniprot-only peptides (Fig. 4e), suggesting that those may derive from erroneous calls or stable peptides from unstable RNAs.

RiboTaper ORFs with peptide support mapped to CCDS genes, with few exceptions (Fig. 4f). We identified peptides belonging to a uORF in the MIEF1 gene^28^ (Fig. 2b)or from dORFs and ncORFs located in conserved and non-conserved genomic regions (Fig. 2c, Supplementary Fig. 12–15). In total, 228 identified peptide sequences were not annotated in Uniprot. In many cases, the novel identified peptides mapped uniquely to their respective ORFs (Fig. 4f, Supplementary Table 4).

## DISCUSSION

Ribo-seq reads of specific lengths can display a precise sub-codon pattern, which allows for accurate identification of the translated frame. However, different experimental protocols can have a marked influence on the sub-codon profiles^2^ (Supplementary Fig. 1). The Ribo-seq protocol is far from being standardized^29^, and codon biases and ribosome stalling^30^ pose challenges to the quantitative understanding of translation at each locus. The RiboTaper method proposed here is robust and tailored to exploit the periodic sub-codon pattern to identify periodic footprint profiles (PFPs) associated with active translation. This principled statistical approach based on the significance of spectra allowed us to independently test transcript regions for PFPs on single loci, yielding high specificity over a wide range of expression and coverage.

Despite the identification of many PFPs in non-coding transcripts^10,11^, evolutionary conservation analysis suggests that the act of translation rather than the production of specific peptides may be the relevant process and may explain the lack of matching peptides^31^. The notable exception was the apparent translation of a considerable number pseudogenes, which may have retained coding potential but are no longer under purifying selection. Quantifying the presence and significance of ribosome footprint reads becomes increasingly difficult for very small translated regions (<20 amino acid long “dwORFs”^7^), and these may be missed by both RiboTaper as well as by conventional mass-spec protocols.

Further investigation is needed to understand the function of ribosomal readout at ncORFs and u/dORFs, considering features of the dynamics of translation (frameshifting, reinitiation), together with other aspects of RNA metabolism which are regulated by ribosomal activity. For example, uORFs can promote transcript susceptibility to RNA-surveillance mechanisms^32^, such as Nonsense-Mediated-Decay. Defining the ensemble of translated uORFs from high-throughput experiments has remained elusive^33^, and our method holds promise in the identification of such events.

Different studies reported a widespread presence of pile-ups at canonical and non-canonical start-codons (NUG) in 5’UTRs^22^. We here considered only AUG start codons, likely missing cases of NUG starting ORFs (Supplementary Fig. 13). Further analyses are needed to investigate the validity of NUGs as efficient start codons. Another possible extension of our current method is the definition of translated alternative RNA isoforms^14,34^. Our approach also does not yet account for frameshifting and may miss cases in which multiple actively translated ORFs overlap with each other.^14^

Recent approaches have used Ribo-seq results to aid peptide discovery^35^. In our hands, the set of identified translated ORFs represented a comparable alternative to public databases and, in fact, a more comprehensive proxy to define the cellular proteome, as the set of RiboTaper PFPs exceeded the coverage of deep mass-spec datasets. Altogether, our approach constitutes a resourceful toolkit for dedicated computational analysis of high-resolution data from Ribo-seq experiments.

## Author Contributions

LC and UO developed the computational approach. LC implemented the RiboTaper method and analyzed the sequencing data, supervised by UO. EW, NM and AH performed the Ribo-seq experiments, supervised by ML. BO carried the evolutionary conservation analysis and helped with the interpretation of the presented findings. HZ, MS and LC analyzed the mass spec data. LC, NM and UO wrote the manuscript, with crucial input from all authors.

## Acknowledgments

LC wants to sincerely thank Roberto Marangoni (University of Pisa) for inspiring and fruitful discussions and Alina Munteanu for help with the Supplementary Figures.

## METHODS

### Ribosome profiling

We followed the original protocol^2^ with minor modifications. For cell lysis, the cell medium was aspirated and cells washed with ice-cold PBS containing 100 μg/ml cycloheximide. No cycloheximide was added to the culture medium before. After thorough removal of the PBS, the plates were immersed in liquid nitrogen and placed on dried ice. For cell lysis, 400 μl mammalian polysome buffer (20 mM Tris-HCl pH 7.4, 150 mM NaCl, 5 mM MgCl2, with 1 mM DTT and 100 μg/ml cycloheximide added freshly) was supplied with 1% (v/v) Triton X-100 and 25 U/ml Turbo DNase (Life Technologies, AM2238) and dripped on the plate which is subsequently placed tilted on wet ice. The cells were scraped off to the lower portion of the dish so that they thawed in lysis buffer. After dispersal of the cells by pipetting, the lysate was triturated ten times through a 26G needle, cleared by centrifugation at 20000g for 5 minutes, flash-frozen in liquid nitrogen and stored at -80 °C until further usage. For isolation of ribosome-protected RNA fragments, 120 μl of the lysate were digested with 3 μl RNase I (Life Technologies, AM2294) for 45 min. at room temperature with rotation. The digestion was stopped by addition of 4 μl Superase-In (Life Technologies, AM2694). Meanwhile, MicroSpin S-400 HR columns (GE Healthcare, 27–5140–01) were equilibrated with 3 ml mammalian polysome buffer by gravity flow and emptied by centrifugation at 600g for four minutes. Of the digested lysate, 100 μl were then immediately loaded on the column and eluted by centrifugation at 600g for two minutes. RNA was extracted from the flow-through, approximately 125 μl, using Trizol LS (Life Technologies, 10296-010). Ribosomal RNA fragments were then removed using the RiboZero Kit (Illumina, MRZH11124) and separated on a 17% denaturing Urea-PAGE gel (National Diagnostics, EC-829). The size range from 27nt to 30nt as defined by loading 20 pmol each Marker-27nt and Marker-30nt was cut out and the RNA fragments subjected to library generation using 3’-Adapter NN-RA3, 5’ adapter OR5-NN, RT primer RTP and PCR primers RP1 (forward primer) and RPI6-7 (reverse primer, containing barcodes). Libraries were sequenced on a HiSeq 2000 device (Illumina). After initial quality control, we obtained ^~^29 Mio raw reads by pooling the RPI6 and RPI7 samples.

Marker-27nt: 5’-AUGUACACGGAGUCGAGCUCAACCCGC-P
Marker-30nt: 5’-AUGUACACGGAGUCGAGCUCAACCCGCAAC-P
NN-RA3: P NNTGGAATTCTCGGGTGCCAAGG-InvdT
OR5-NN: 5’-GUUCAGAGUUCUACAGUCCGACGAUCNN
RTP 5’-GCCTTGGCACCCGAGAATTCCA
RP1 5’-AATGATACGGCGACCACCGAGATCTACACGTTCAGAGTTCTACAGTCCGA
RPI6 5’-CAAGCAGAAGACGGCATACGAGATATTGGCGTGACTGGAGTTCCTTGGCACCCGAGAATTCCA
RPI7 5’-CAAGCAGAAGACGGCATACGAGATGATCTGGTGACTGGAGTTCCTTGGCACCCGAGAATTCCA

### Preprocessing of Ribo-seq and RNA-seq

Poly-A selected RNA-seq data for HEK293 was obtained from a recent study^36^ (accession: GSM1306496). Ribo-seq reads were stripped from the adapter sequences, and reads aligning to rRNA sequences were discarded using Bowtie^37^. Unaligned Ribo-seq reads and RNA-seq reads were aligned to the human genome (hg 19) using the split-aware aligner STAR^38^. The STAR genome index was built using annotation obtained from GENCODE (version 19)^39^. For RNA-seq and Ribo-seq, a maximum of 4 mismatches were allowed and multi-mapping to up to 8 different position was permitted. Alignments flagged as secondary alignment were filtered out, ensuring 1 genomic position per aligned read. Ultimately, we thus obtained ^~^25 mio aligned reads. To infer the P-site locations, we created a histogram of distances between the 5’end of Ribo-seq reads and well-annotated start and stop codons (CCDS, see below), for each read length (Fig. 1a, Supplementary Fig. 1). Read lengths and offsets used to infer the P-sites position are available in the Supplementary Table 1 for the different Ribo-seq libraries used. RNA-sites were calculated using an arbitrary (26th) position for each RNA-seq read. TPM values for RNA-seq and Ribo-seq were calculated using RSEM^40^. Custom Unix scripts and BedTools^41^ were used to create data tracks containing 1) Ribo-seq coverage; 2) P-sites distributions, 3) RNA-seq coverage and 4) RNA-sites distributions for different genomic regions.

### Exon-level annotation, simulations and ORF identification

Data tracks were created for each annotated exon in the GENCODE v19 annotation, distinguishing between regions annotated as consensus coding sequences (CCDS), non-CCDS exons inside CCDS-containing genes, and exons in non-CCDS genes. Non-CCDS regions (5’UTRs, alternative exons etc…) were annotated with respect to CCDS locations. Exons with more than 5 P-sites were considered for quality control checks. For the benchmarking test, we sampled 1000 CCDS exons from different read lengths and coverage as a positive set. For each exon, we randomly shuffled 1000 times the P-sites positions to obtain a negative set.

Full ORFs were detected by merging exons according to the transcript structures reported in the GENCODE v19 annotation. For CCDS genes, all CCDS transcripts comprising the “appris” tag were used, by prioritizing transcript with the “appris_principal” tag. For non-CCDS transcripts, all annotated transcripts containing an exon with >5 P-sites were used. For Zebrafish, all transcripts structures annotated in Ensembl (version 76) were used.

For each transcript, every pair of consecutive start-stop codons (ORF) was tested for its 3nt periodic pattern using the multitaper method, in all the three possible frames (p-value <0.05). ORFs with less than 50% of in-frame P-sites were excluded. In case of multiple possible start codons, we chose the most upstream in-frame ATG with more than 5 P-sites positions (>50% in-frame) between it and its closest neighbor ATG (Fig. 2a). In case of multiple transcript isoforms harboring the same ORF, the transcript with the best support from RNA-seq was chosen.

ORFs were annotated as follows: ORFs_ccds as ORFs overlapping known CDS regions in CCDS genes; non-CCDS coding ORFs were defined similarly, but when overlapping non-CCDS CDS regions. uORFs and dORFs as ORFs in CCDS genes non overlapping with any CDS exon, and annotated with respect to the annotated transcript CDS; ncORFs as ORFs in non-CCDS genes and not overlapping with any CDS exon (Supplementary Fig. 7). ORFs were filtered out when >30% of the Ribo-seq coverage supported by multi-mapping reads only (filtering was disabled for the custom peptide database creation).

Multitaper method

In digital signal processing, the quantitative estimation of periodic components in a finite signal (the power spectral density, or PSD) is an intense area of study, with application to diverse fields of scientific research. A switch from the original representation of a signal (the time domain) to its spectrum of fixed periodic components (the frequency domain) is achieved via the Fourier Transform. In the frequency domain, a vector of coefficients represents the contribution of each frequency component in shaping the original signal.

The raw output of the Fourier transform (the periodogram) typically suffers from high variability, and as such, it represents a poor estimate of the PSD. Moreover, the limited amount of available realizations of the same signal (i.e. the lack of replicates) poses a real challenge in the estimation of robust coefficients for different frequencies. Applying a smoothing window (taper) to the signal before calculating its Fourier transform helps reducing such variability, but generally creates a biased estimate of the PSD. The choice of taper functions is fundamental, and many different solutions have been proposed in the last decades to find the “optimal” window function.

The multitaper method, originally proposed by Thomson^18^, offers a promising, non-parametric solution to the PSD estimation problem. Its central idea is to apply a set of multiple window functions to the signal, and average the spectral estimates of the ensemble of tapered signals. As the window functions used in the multitaper method are a set of orthonormal functions, the resulting spectra are independent and can be averaged. Specifically, discrete prolate spheroidal sequences (dpss, or Slepian sequences) have been shown to maximize the information content of a finite signal at a given frequency resolution. The use of the Slepian sequences in the multitaper method allows for an optimal solution for reducing the variance in the power spectrum.

The use of multiple orthonormal window functions on the same signal also enables us to test the robustness of the estimated spectral coefficients. The amount of variance captured by the estimated coefficient at each frequency bin can be compared against a null hypothesis of white noise, leading to a reliable statistical test to determine significant frequency components^20^.

### RiboTaper

The original multitaper algorithm from Thomson is implemented in R in the package “multitaper”^42^. A stretch of zeros was added to the input sample to reach a minimum length of 50 nt. The multitaper was run with 24 tapers, setting the time-window parameter to 12. Moreover, sequences shorter than 500 nt were zero-padded to 1024 data points before computing the Discrete Fourier Transform, to obtain an adequate frequency resolution in the spectrum. F-values were extracted from the frequency bin closest to 3nt periodicity. P-values from the F-statistic^20^ were calculated by using 2 and 2k-2 degrees of freedom, where k is the number of tapers (24 in this study). ORFs and exons with less than 6 P-sites or shorter than 6 nt were ignored.

### QTI-seq comparison

For every reported QTI-seq peak^22^, we selected the closest ORF called by RiboTaper based on the reported distance relative to the annotated start codon. Only ATG start codons were used.

### Conservation Analysis

PhastCons scores were extracted as average over the entire ORF, or in 25nt windows around start and stop codon. ORFs were then scored with PhyloCSF in the “mle” (default) mode, using the “29mammals” parameter set on the 46-vertebrate alignment to the human genome (hg19) after alignment filtering steps as described in Bazzini *et al.*^6^ We additionally used the hexamer score from the CPAT tool to assess the coding potential of different ORFs, using the available trained model for the human genome. For each category, the scores were compared against a control set of ORFs matching length and conservation of the category of interest. For ORFs_ccds and nonccds coding ORFs, we selected ORFs shorter than 300nt as meaningful matching controls (Supplementary Fig. 10). SNPs were downloaded as .gvf files from Ensembl (v75, 1000 Genomes phase 1). We removed SNPs in reverse orientation, SNPs falling into genomic repeats (using the RepeatMasker track from the UCSC genome browser, March 22, 2015), and rare SNPs with derived allele frequency <1%. We then recorded for each ORF and its conceptual translation the number of synonymous and nonsynonymous SNPs when comparing to the human reference genome, as well as the number of synonymous and nonsynonymous sites derived from the degeneracy of the genetic code. For every set of ORFs, we aggregated these numbers and calculated the dN/dS ratio, where dN is the number of nonsynonymous SNPs per nonsynonymous site, and dS the number of synonymous SNPs per synonymous site, respectively. For the CPAT and PhyloCSF scores, p-values come from Wilcoxon-Mann-Whitney tests. For dN/dS ratios, p-values come from a Chi-square test, using as expected frequencies the values from the ORF control set^43^.

### Mass-spec sample collection and preparation

The proteomic data for HEK293 was published recently^27^ (PRIDE accession number: PXD002389). Briefly, cells were grown in DMEM (life technologies). Lysis was performed in 50 mM ammonium bicarbonate buffer (ABC, pH 8.0) containing 2% SDS and 0.1 M DTT. Sulfhydryl groups were alkylated by adding iodoacetamide to a final concentration of 0.25 M and incubation for 20 min. Proteins were precipitated, resuspended in 6M urea/ 2M thiourea/ 10mM HEPES and digested into peptides using LysC (3 h) and Trypsin (overnight, diluted 4x with 50 mM ABC). Peptides were then acidified, desalted and subjected to isoelectric focusing (IEF) for fractionation.

### LC-MS/MS and data analysis

Peptides were desalted using stage tip purification and subsequently analyzed by online liquid-chromatography tandem mass-spectrometry on a Q-Exactive (ThermoFisher) instrument using nano-electrospray ionization. Resolution was set to 70,000 and 17,500 for full and fragments scans respectively. Peptides were identified from MS/MS spectra by searching against the recent Uniprot human database (2014-10) or the newly generated HEK293 specific database using ribosome profiling using MaxQuant^44^ version 1.5.2.8. For all searches carbamidomethyl (C) was set as fixed, oxidation (M) and acetylation (protein N-term) and deamidation (NQ) as variable modifications. A maximum of two missed cleavages was allowed. Peptide FDR was set to 0.01, minimal peptide length was set to 7 amino acids and the main search peptide tolerance was set to 4.5 ppm. The protein FDR was disabled.

### Mass-spec data processing

Custom peptide databases were built by using all the set of identified ORFs prior to filtering for multi-mapping reads. FDR was calculated based on the ratio of hits in the positive and decoy database, as previously described^44^. Counts and feature distribution (PEP, Score, Sequence length) from evidence files were compared based on a FDR < 1 %, excluding Reverse Hits and Contaminants as well as using unique sequence information. Non-Uniprot peptide sequences were defined using PeptideMatch^45^.

### Accession codes

Gene Expression Omnibus (GEO) GSE73136 (Ribo-seq data from HEK293 cells).

### Software availability

All the scripts (written in Unix, Bedtools and R) to create the data tracks, run the multitaper analysis, annotate the exons, perform the simulations, reconstruct the full ORFs etc. are available at https://ohlerlab.mdc-berlin.de/software and in the Supplementary Software. The method requires a genome fasta file, a GTF file for annotation, alignment files for Ribo-seq and RNA-seq and the read lengths (with respective cutoffs) used for the P-sites calculation.

Supplementary Figure 1: Metagene analysis for different datasets. Aggregate plots in different read lengths (from 25 to 30 nt) are shown, showing distance between 5’ends and annotated start and stop codons. Distinct profiles, in terms of both precision and coverage, emerge in the different datasets. a) HEK293 - This study; b) HEK293, Gao *et al* 2014; c) Zebrafish 5h post-fertilization, Bazzini *et al* 2014.

Supplementary Figure 2: Quality control steps in the different datasets used. HEK293, this study: a) Ribo-seq and b) RNA-seq coverage of the different used datasets, together with the number of defined P-site positions. c) Number of CCDS exons with more than 5 Ribo-seq and RNA-seq reads. d) % of CCDS exons called by RiboTaper or Chi-squared test using Ribo-seq (p-value lower than 0.05), together with % of negative exons using RNA-seq (p-value higher than 0.05). e) Histogram of p-values for the multitaper and f) Chi-squared test on CCDS exonic RNA profiles. g) Accumulation of P-sites on the maximum frame for exons in different transcripts regions. h) Frame agreement with the CCDS annotation. j) % of periodic exons in different length-coverage categories, derived by taking the second, fourth and sixth of seven quantiles of length-coverage distributions, using Ribo-seq or j) RNA-seq.

Supplementary Figure 3: Quality control steps in the different datasets used. HEK293, Gao et al: a) Ribo-seq and b) RNA-seq coverage of the different used datasets, together with the number of defined P-site positions. c) Number of CCDS exons with more than 5 Ribo-seq and RNA-seq reads. d) % of CCDS exons called by RiboTaper or Chi-squared test using Ribo-seq (p-value lower than 0.05), together with % of negative exons using RNA-seq (p-value higher than 0.05). e) Histogram of p-values for the multitaper and f) Chi-squared test on CCDS exonic RNA profiles. g) Accumulation of P-sites on the maximum frame for exons in different transcripts regions. h) Frame agreement with the CCDS annotation. j) % of periodic exons in different length-coverage categories, derived by taking the second, fourth and sixth of seven quantiles of length-coverage distributions, using Ribo-seq or j) RNA-seq.

Supplementary Figure 4: Quality control steps in the different datasets used. Zebrafish, Bazzini et al: a) Ribo-seq and b) RNA-seq coverage of the different used datasets, together with the number of defined P-site positions. c) Number of CDS exons with more than 5 Ribo-seq and RNA-seq reads. d) % of CDS exons called by RiboTaper or Chi-squared test using Ribo-seq (p-value lower than 0.05), together with % of negative exons using RNA-seq (p-value higher than 0.05). e) Histogram of p-values for the multitaper and f) Chi-squared test on CDS exonic RNA profiles. g) Supplementary Figure 6: Accumulation of P-sites on the maximum frame for exons in different transcripts regions. h) Frame agreement with the CDS annotation. j) % of periodic exons in different length-coverage categories, derived by taking the second, fourth and sixth of seven quantiles of length-coverage distributions, using Ribo-seq or j) RNA-seq.

Supplementary Figure 5: Multitaper performance for different numbers of tapers. Shown are the a) AUC, b) sensitivity and c) specificity values for real and shuffled exonic P-sites profiles, along different length-coverage categories, derived by taking the second, fourth and sixth of seven quantiles of length-coverage distributions. Little change is detectable by using more than 24 tapers.

Supplementary Figure 6: Different methods performances on P-sites and RNA-sites tracks for coding exons of different length/coverage values. Shown are different ORFscore cutoffs, Chi-squared and multitaper statistical tests (see text for more details). a) Data in HEK 293 cells from our study (CCDS annotation); b) Data in Zebrafish 5hpf from Bazzini et al, 2014.

Supplementary Figure 7: Schematics of RiboTaper ORF annotation. In a) a “uORF” is defined as upstream of the annotated start codon and non-overlapping any coding exon, while different “ORFs_ccds” are overlapping annotated coding exons. A “dORF” is defined as downstream of the stop codon and not overlapping any coding exon. Shown is also a lincRNA ORF overlapping a coding exon, therefore annotated as “nonccds_coding_ORF". In b) a “nonccds_coding_ORF” in a non-CCDS protein coding gene, defined as overlapping a coding exon. A “nonccds_coding_ORF” in a processed_transcript gene is also present. An ncORF is defined as an ORF in a non-CCDS gene not overlapping any coding exon, here in an antisense gene. In c) an ncORF in a lincRNA gene is shown.

Supplementary Figure 8: FLOSS scores for ORFs identified by RiboTaper. Shown are FLOSS scores with their cumulative distributions for a) CCDS genes and ORFs_ccds. b) 5’UTRs and uORFs, c) 3’UTRs and dORFs, d) non-coding genes and different ncORFs categories. FLOSS values and cutoffs were calculated as in Ingolia et al, 2014. Low FLOSS scores indicate a protein-coding like fragment length distribution.

Supplementary Figure 9: The toddler ncORF. Shown are P-sites position, RNA-seq coverage and ORF position. Data from Bazzini et al, 2014.

Supplementary Figure 10: Length and conservation-matched ORFs for the conservation analysis. No significant difference in terms of a) length and b) nucleotide conservation (PhastCons) was found between detected ORFs and randomly chosen controls ORFs. c) Scores from PhyloCSF (used in the -mle mode) agree with the ones from CPAT (see main text and Figure 4 for more details). * = p-value<0.05; **= p-value<0.01; ***= p-value<0.001, Wilcoxon-Mann-Whitney test. ORFs_ccds and nonccds coding ORFs were selected if <300 nucleotides long, to match negative controls ORFs.

Supplementary Figure 11: Additional statistics about Ribotaper- and Uniprot-only identified peptides. a) Overlap between genes with peptide evidence in the different search strategies. b) Overlap of Peptide Spectrum Matches (PSMs) in the two strategies. c) FDR vs. PSMs count for the two search strategies. d) Comparisons of RiboTaper-only identified peptides vs. Uniprot-only identified peptides (PEP=Posterior Error Probability).

Supplementary Figure 12: Genomic locations of “non-canonical” ORFs with peptide evidence. A dORF in the SMIM20 gene. Below a screenshot from the GWIPS-viz Genome Browser.

Supplementary Figure 13: Genomic locations of “non-canonical” ORFs with peptide evidence. A lincRNA ncORF in the LOC93622 gene. Below a screenshot from the GWIPS-viz Genome Browser. A CUG-start codon is supported by initiating ribosomes from QTI-seq and LTM data. The second exon (on the right) showing a strong accumulation of RNA-seq reads is present in circBase, http://www.circbase.org/, (hsa_circ_0069092).

Supplementary Figure 14: Genomic locations of “non-canonical” ORFs with peptide evidence. An antisense ncORF in the CTD-2162K18.5 gene, in a non-conserved genomic region. Below a screenshot from the GWIPS-viz Genome Browser.

Supplementary Figure 15: Genomic locations of “non-canonical” ORFs with peptide evidence. An antisense ncORF in the RP11–139H15.1 gene (on the left), in a conserved genomic region. Below a screenshot from the GWIPS-viz Genome Browser.

Supplementary Table 1: Statistics about alignment and pre-processing of the sequencing datasets used in this study.

Supplementary Table 2: Complete list of identified ORFs in the different datasets used.

Supplementary Table 3: Filtered list of identified ORFs in the different datasets used. Non-coding ORFs overlapping CDS regions were removed. Additionally, ORFs were filtered out when >30% of the Ribo-seq coverage were supported by multi-mapping reads only.

Supplementary Table 4: Evidence table for peptides identified by the RiboTaper database only.

Supplementary File 1: Archive containing bed files for the identified ORFs.

Supplementary Software: RiboTaper (version 1.0) software code.a)

